# The Global Phosphorylation Landscape of Acute Heat Stress in Reef-Building Corals

**DOI:** 10.64898/2026.10.29.762943

**Authors:** Immy A. Ashley, Giada Tortorelli, Jasmine Nguyen, Kareem Alba, Eva Majerová, Declan M. Winters, Shaylyn Carthen, Tahirih Perez, Sabrina L. Rosset, Crawford Drury, Mehdi Bouhaddou

**Affiliations:** Department of Microbiology, Immunology, and Molecular Genetics; University of California, Los Angeles, Los Angeles, CA, USA; Institute for Quantitative and Computational Biosciences; University of California, Los Angeles, Los Angeles, CA, USA; Molecular Biology Institute, University of California Los Angeles; Los Angeles, CA, USA; Hawaiʻi Institute of Marine Biology; University of Hawaiʻi at Mānoa, Kāneʻohe, HI, United States

## Abstract

Warming oceans disrupt the coral-algal symbiosis that sustains reefs, yet the signaling mechanisms initiating bleaching remain poorly understood. Integrating global phosphoproteomics across three reef-building corals with protein structure modeling revealed conservation between coral and human kinases, enabling inference of coral kinase activities and uncovering a shared signaling program induced by acute heat stress. Heat rapidly activated ERK, JNK, and p38 MAP kinases, suppressed cell cycle signaling, and remodeled pathways controlling chromatin, RNA processing, and metabolism. Most strikingly, thermal stress rewired immune receptor signaling networks associated with non-self recognition, suggesting that heat shifts the coral host from tolerance to immune recognition of its algal symbionts. These findings define conserved signaling mechanisms that precede bleaching, revealing actionable molecular targets for enhancing coral resilience.

## INTRODUCTION

Coral reef ecosystems are under increasing pressure due to a range of stressors, including anthropogenic climate change (*1*). Warming oceans cause coral bleaching, the breakdown of the symbiosis between the coral and their intracellular photosynthetic dinoflagellate algae, leading to compromised function and eventual mortality. These events are becoming more frequent and severe (*2*) and global oceans are committed to several degrees of warming even if emissions stopped today (*3*). The symbiosis between scleractinian corals and their intracellular dinoflagellate algae (*Symbiodiniaceae*) is the energetic foundation that supports the structure and function of coral reefs (*4*). This evolutionarily ancient relationship is the key determinant of the persistence of corals across geologic time (*5*), but is particularly vulnerable to changing environmental conditions (*6*). Thermal stress can disrupt this relationship, leading to coral bleaching, yet the molecular mechanisms initiating the symbiosis breakdown remain poorly understood (*7–9*).

Several, non-mutually exclusive mechanisms have been proposed to drive this symbiosis breakdown. Imbalances in nutrient exchange can destabilize metabolic homeostasis between host and symbiont, while the build-up of reactive oxygen species arising from the host and/or symbiont can overwhelm cellular antioxidant defenses and cause oxidative stress (*10–17*). These stresses have been associated with diverse cellular responses, including the unfolded protein response, DNA damage, cell cycle suppression, innate immunity, programmed cell death, and ultimately the loss of the algal symbiont (*10*, *18*). However, it remains unclear which responses represent upstream signaling events and which reflect downstream consequences. The early signaling events triggered by heat stress remain poorly characterized, making phosphorylation an especially informative readout because it represents one of the most rapid cellular responses to external stimuli.

Organisms must sense and respond to their environment rapidly and efficiently, supporting homeostasis and survival, especially in response to harmful conditions (*19*). Phosphorylation, a post-translational modification catalyzed by protein kinases, is a central regulator of protein function, capable of altering protein activity, structure, localization, and molecular interactions, and thereby serves as a major mechanism by which cells control biological phenotypes. Furthermore, phosphorylation is central to cell biology, with estimates in humans indicating that 75% of the proteome can be phosphorylated, highlighting the ubiquity of this modification (*20*). Here, we sought to define how kinase-driven signaling networks in corals are remodeled early during heat stress and theorize how these changes could contribute to the initiation of symbiosis breakdown.

The use of ‘omics’ techniques creates an opportunity to advance the cell biology of reef-building corals towards the standard of the human medical field, bridging the gap between model and non-model systems (*21*, *22*). Healthy and disrupted symbioses have been studied through the lenses of transcriptomics, genomics, proteomics and metabolomics in response to a wide range of environmental stressors, including heat, ocean acidification, light, salinity, and nutrients (*23–31*). However, phosphoproteomics, which can measure changes in tens of thousands of phosphorylation sites simultaneously, has not yet been reported in reef-building corals and has only been applied to Aiptasia, a model sea anemone (*32*). Although vertebrate phosphorylation networks have been extensively characterized, comparable knowledge in corals remains limited, representing a major gap in the understanding of how heat stress initiates symbiosis breakdown and bleaching.

To address this gap, we developed a mass spectrometry phosphoproteomics approach that enables the direct measurement of protein phosphorylation in corals in response to acute heat stress. We first optimized a sample preparation pipeline for state-of-the-art phosphopeptide recovery from reef-building corals. We then modeled the structures of coral kinases using AlphaFold2 and quantitatively compared their structural and sequence similarities with human orthologs. Leveraging this conservation, we applied human kinase-substrate motif information to the coral phosphoproteome to infer changes in kinase and pathway activities following acute heat stress. Together, our results reveal a conserved kinase landscape and heat stress signaling program across reef-building corals, identifying molecular pathways that may represent targets for enhancing coral resilience to warming oceans.

## RESULTS

### Acute thermal stress drives rapid remodeling of the coral phosphoproteome

We performed global mass spectrometry abundance proteomics and phosphoproteomics to elucidate the coral phosphorylation landscape during acute thermal stress (Fig. 1A). Three reef-building coral species—*Pocillopora acuta*, *Montipora capitata*, and *Porites compressa*—were subjected to short-term exposure to 32 ℃, with a control set maintained at ambient temperature. This temperature is beyond the typical temperature range of our study system (Kaneʻohe Bay, Oʻahu, Hawaiʻi) and is traditionally used in experiments to elucidate rapid heat-stress signaling and to trigger heat stress-response pathways (*33*). Samples were collected before heat treatment and 1-, 2-, and 4-hours after the onset of heat exposure, processed to split the host and symbiont fractions and subjected to paired mass spectrometry abundance proteomics and phosphoproteomics to quantify total protein levels and phosphorylation site abundances, respectively. Abundance proteomics analysis detected a total of 9608 unique proteins in *P. acuta,* 6898 in *M. capitata*, and 7298 in *P. compressa* (Fig. 1B, fig. S1). Using an untargeted phosphoproteomics approach, we were able to identify a total of 13455 unique phosphopeptides in *P. acuta,* 6586 in *M. capitata*, and 5341 in *P. compressa* (Fig. 1C). Importantly, peptide intensities were strongly correlated between biological replicates (Pearson’s r >0.7), demonstrating robust experimental reproducibility (Fig. 1D). As expected, changes in protein level phosphorylation were initiated earlier and were greater in number of changes relative to changes in protein levels (Fig. 1E-G), with 26-36% of phosphosites significantly altered in at least one time point across species compared to only 4-20% of proteins.

**Figure 1:**
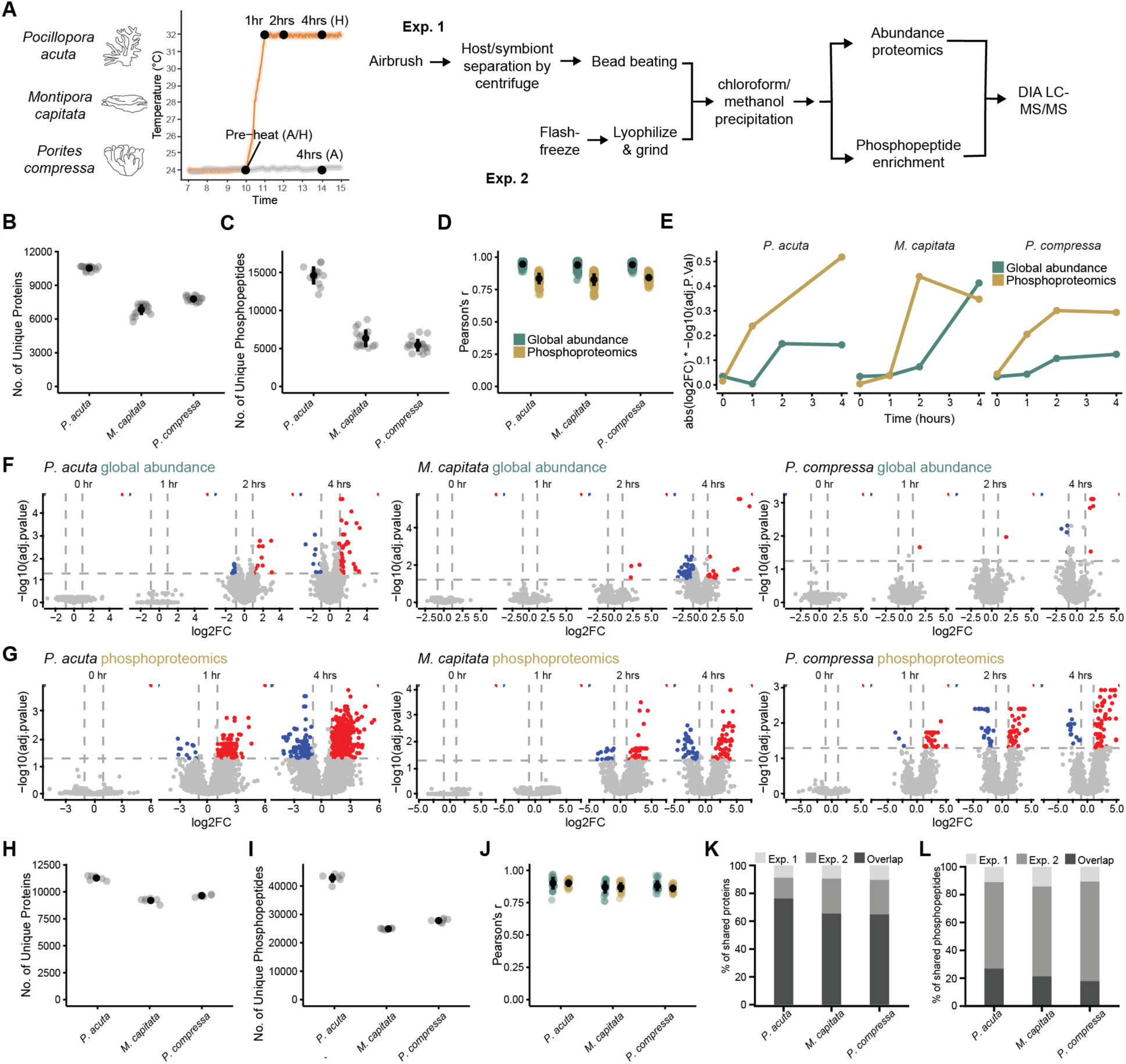
Acute heat stress remodels the coral phosphoproteome despite minimal changes in protein abundance. **(A)** Experimental schematic of the species used (*Pocillopora acuta*, *Montipora capitata, Porites compressa*), temperature conditions, data collection and processing for proteomics and phosphoproteomics analysis; **(B)** Unique proteins identified in the time-course experiment across the three species; **(C)** Unique phosphopeptides identified in the time-course experiment for the three species; **(D)** Pearson’s r correlation of the global proteomics and phosphoproteomics data within time points, across the experiment; **(E)** Absolute log fold-change of global proteomics and phosphoproteomics data across the experiment; **(F)** Volcano plots of the global proteomics changes across time; **(G)** Volcano plots of phosphoproteomics changes across time; **(H)** Unique proteins identified in the validation experiment across the three species; (**I)** Unique phosphopeptides identified in the validation experiment for the three species; **(J)** Pearson’s r correlation of the global proteomics and phosphoproteomics data across the experiment; **(K)** Percentage of proteins detected across the two experiments (proteins unique to experiment 1 (998, 875, and 1008) and experiment 2 (1684, 2308, and 2408); **(L)** Percentage of phosphopeptides detected across the two experiments (phosphopeptides unique to experiment 1 (5052, 3903, and 3130) and experiment 2 (28291, 17667, and 20891).

To validate our results, we repeated the experiment with sampling at the 4-hour time point only. In this experiment, fragments of coral were immediately flash frozen, lyophilized, then ground with a pestle and mortar without separating host and symbionts to minimize the impacts of the initial sample processing on the phosphorylation signaling dynamics (Fig. 1A). Our modified sample processing protocol markedly improved our identifications, increasing protein detections by 17-20% and phosphopeptide detections by 160-320% relative to experiment 1. In experiment 2 we identified 11292 unique proteins in *P. acuta*, 8331 in *M. capitata* and 8698 in *P. compressa* (Fig. 1H). Additionally, we detected 35129 unique phosphopeptides in *P. acuta*, 19576 in *M. capitata* and 22510 in *P. compressa* (Fig. 1I). Correlation within replicates at each time point was similarly high (Fig. 1J). Across the two experiments we detected 8610, 6023, and 6290 shared proteins in *P. acuta*, *M. capitata* and *P. compressa*, respectively; corresponding to 76.2%, 65.4% and 64.8% of the combined data (Fig. 1K). We detected 12231, 5817, and 5213 shared phosphopeptides in *P. acuta*, *M. capitata,* and *P. compressa* across the two experiments respectively, corresponding to 26.8%, 21.2%, and 17.8% of the combined data.

### Protein kinases are structurally conserved between corals and humans despite sequence divergence

Since kinase substrate specificities have not been defined in corals, we sought to leverage human-derived specificity profiles to infer kinase activity from our phosphoproteomics data. We first assessed the evolutionary conservation of coral and human kinases using OrthoFinder to match coral proteins to their human orthologs (S. table 1 and 2) (*34*) and AlphaFold2 to predict structural similarities between human and coral kinases (Fig. 2A, S. table 3). Using OrthoFinder, we identified coral orthologs for 53.7% (*P. acuta*), 53.3% (*M. capitata*), and 54.4% (*P. compressa*) of all human genes, genome wide (Fig. 2B). Restricting this analysis to human kinases alone increased the match rate to 70.3%, 70.9%, and 71.7% across species, respectively, indicating that kinases are more conserved than the genome average. When we further required the matched coral gene to contain a recognized kinase domain, the match rate remained high at 64.2%, 64.8%, and 65.4%, respectively (Fig. 2B). Together, these findings demonstrate substantial conservation of kinases between humans and corals despite their deep evolutionary divergence.

**Figure 2:**
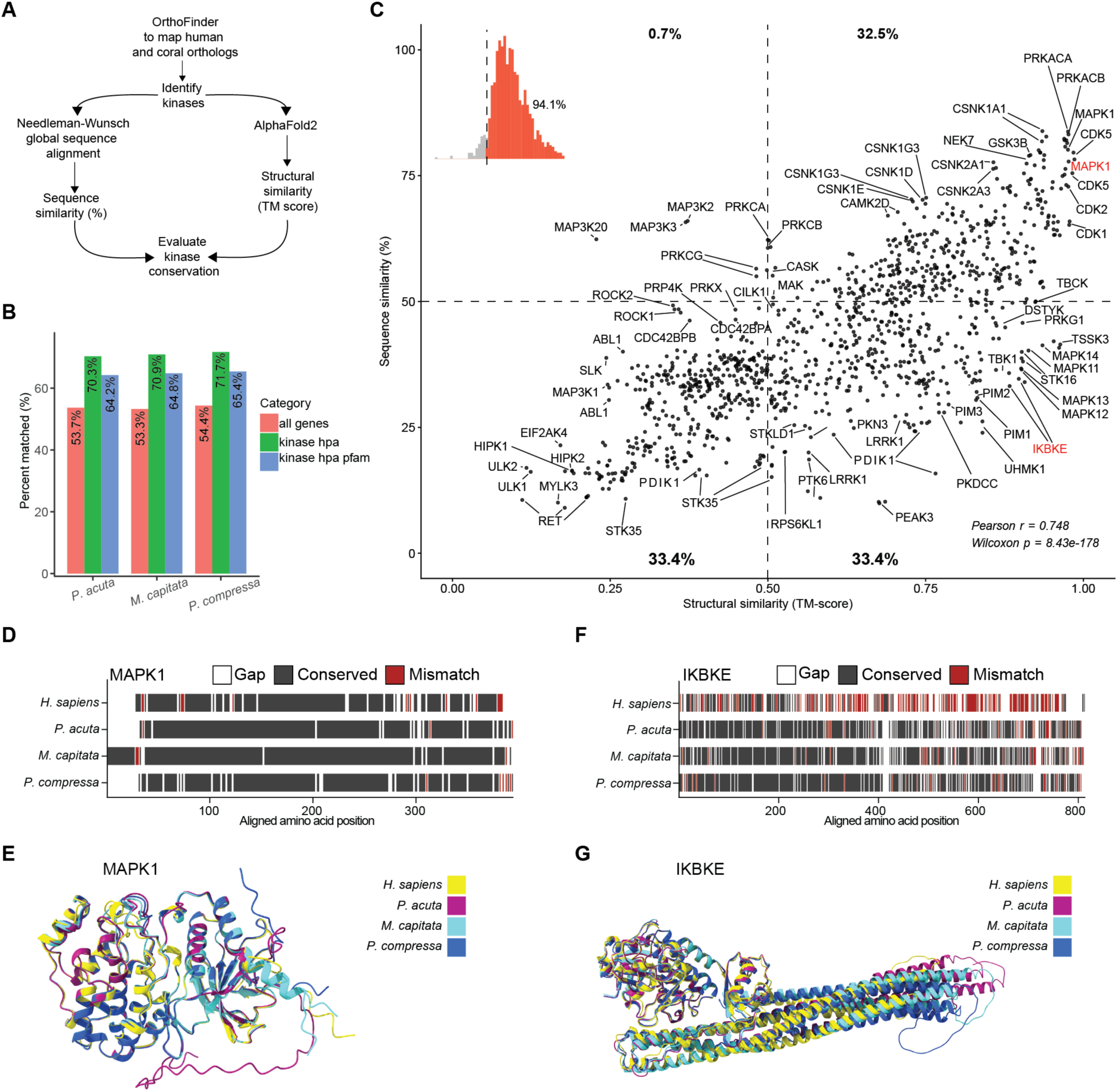
Coral and human kinases are structurally conserved. **(A)** Bioinformatic schematic of the identification of putative coral kinases using homology to human kinases; **(B)** Percentage matches of all human genes identified across *M. capitata, P. acuta and P. compressa,* matches of all human kinases from the human protein atlas (HPA), and all human kinases from the human protein atlas (HPA) for which the matching coral ortholog has an active kinase domain; **(C)** The correlation of structural and sequence similarity of identified putative coral kinases, with both a match to a human kinase, and an active kinase domain. Inset: distribution of the per-kinase difference between structural similarity (TM-score) and scaled sequence similarity (Sequence Similarity/100); positive values (red) indicate kinases for which structural similarity exceeds sequence similarity; **(D)** The sequence alignment of a representative protein (putative coral MAPK1) with a high sequence and high structural similarity; **(E)** The structural alignment of putative coral MAPK1; **(F)** The sequence alignment of a representative protein (putative coral IKBKE) with a low sequence and high structural similarity; **(G)** The structural alignment of putative coral IKBKE.

We used all coral genes that matched a human kinase and contained a kinase domain for comparative sequence and structure analyses (Fig. 2C, fig. S2, S. data 1). As expected, sequence (% identity) and structural (TM-score) similarity were positively correlated (Pearson’s r = 0.748, Wilcoxon *p* = 8.43 × 10⁻¹⁷⁸); however, numerous exceptions to this relationship were observed. Interestingly, although only 33.3% of matched kinases had a sequence identity above 50%, 65.9% had a TM-score above 0.5, indicating substantial conservation of protein structure despite divergence at the sequence level (Fig. 2C). Although 32.5% of kinases exhibited high sequence and structural similarity, such as MAPK1 (Fig. 2D-E), 33.4% of kinases exhibited high structural similarity despite low sequence similarity, such as IKBKE (Fig. 2F-G). In contrast, only 0.7% exhibited high sequence similarity but low structural similarity. Beyond these threshold-based categories, a per-kinase comparison of relative conservation revealed that structural similarity exceeded sequence similarity in 94.1% of matched kinases (Fig. 2C, inset), indicating that the tendency for structure to be retained beyond sequence identity is not confined to kinases meeting an absolute conservation threshold, but is a near-universal feature of the coral kinome. This structural conservation further supports the possibility that the functional roles of these kinases are also conserved, providing a rationale for applying human prior knowledge networks to infer kinase activities in coral.

### Thermal stress activates conserved MAPK signaling in reef-building corals

We next sought to infer how kinase signaling activities were remodeled by heat stress. Conventional kinase activity analyses using phosphoproteomics data integrate prior knowledge networks of experimentally defined kinase-substrate interactions with changes in substrate phosphorylation to estimate kinase activity (*35*, *36*). Such networks are not available for corals, and substantial sequence divergence between coral and human proteins complicates direct transfer phosphorylation site locations between coral and human proteins. We therefore adopted a motif-based approach using The Kinase Library algorithm, which predicts kinase-substrate relationships from the amino acid sequence surrounding each phosphosite (*37*, *38*), permitting direct application to coral protein sequences. Although the algorithm was developed from data derived from recombinant human kinases, we reasoned that human kinase-substrate motif specificity information could be transferable to corals given the substantial structural conservation between human and coral kinases (Fig. 2). The Kinase Library differential phosphorylation analysis infers changes in kinase activities by assessing coordinated changes in the abundance of phosphorylation sites exhibiting kinase-responsive consensus sequence motifs. Predicted serine/threonine kinase activities that were significantly (adjusted p < 0.05) up- or down-regulated in at least one species at 4 hours post heat stress were grouped into 10 functional classes (Fig. 3A, S. table 4), confirmed by pathway enrichment analysis (Fig. 3B).

**Figure 3:**
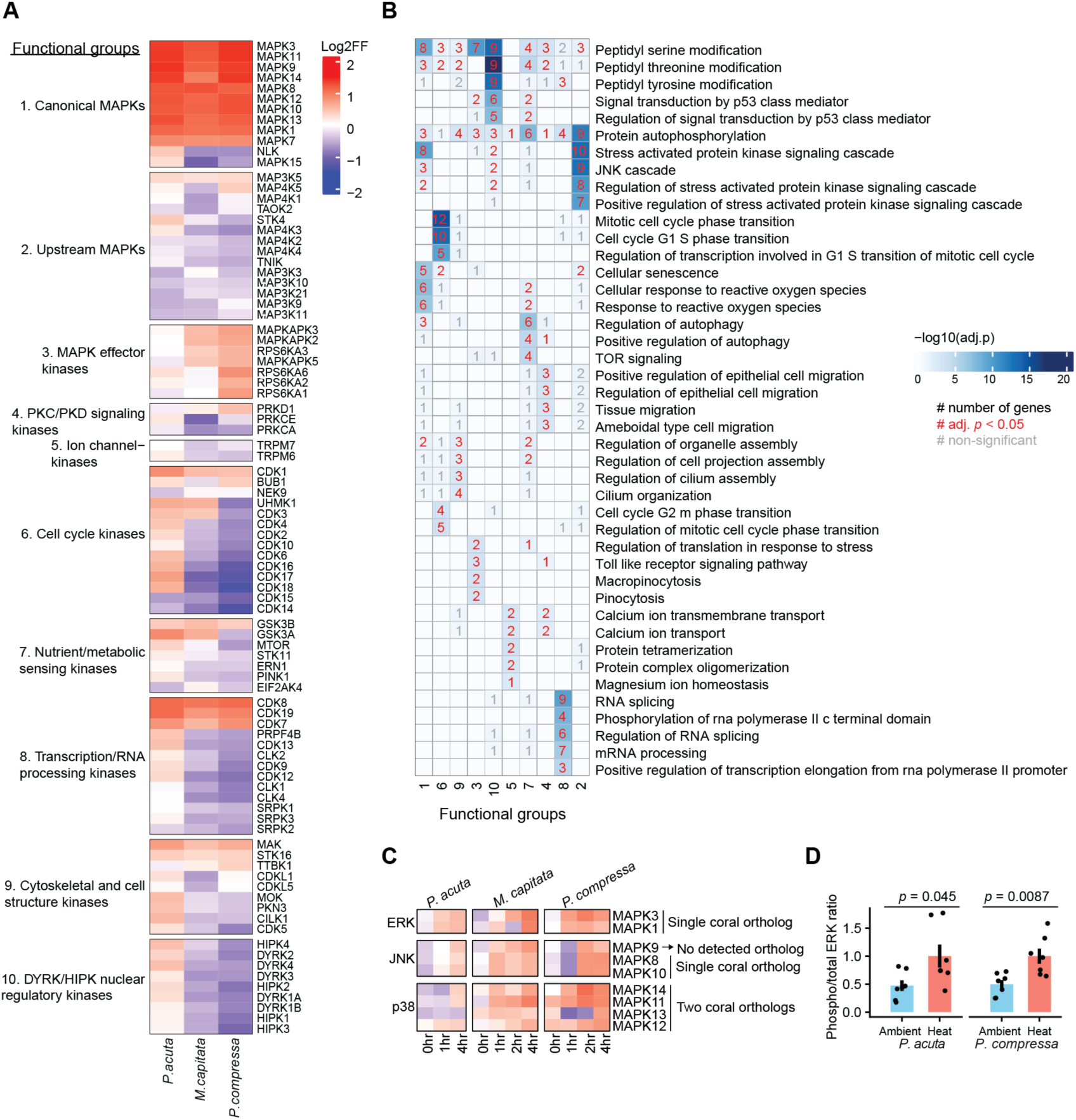
Thermal stress remodels kinase activities across a broad range of functional classes. **(A)** Predicted kinase activity at 4h after thermal stress, clustered using k-means, from validation experiment (Exp 2); **(B)** Enriched pathways in significantly regulated kinases, grouped by classes in A. Number (#) indicates the number of genes detected within that term, red indicating the term is significantly enriched whereas grey indicates not significant. No number indicates no genes detected in that term from that functional group; **(C)** Time dynamics of ERK, JNK, and p38 pathway activities, with noted presence/absence of identified coral orthologs from time course experiment (Exp 1); **(D)** Ratio of total-ERK to phosphorylated-ERK in *P. acuta* and *P. compressa* measured by ELISA

The canonical MAPK cluster showed a strong and conserved response, with all members of the ERK (MAPK1, MAPK3), JNK (MAPK8-10), and p38 kinases (MAPK11-14) strongly activated across all three species. Interestingly, upstream MAPKs were predominantly suppressed across species, whereas downstream MAPK effector kinases tended towards activation in *M. capitata* and *P. compressa*, while *P. acuta* showed a comparatively muted response. Cell cycle, transcription, RNA processing, cytoskeletal, and DYRK/HIPK nuclear regulatory kinases trended toward strong suppression in *M. capitata* and *P. compressa*, diverging from *P. acuta*, which trended toward activation. Exceptions to this pattern included the transcription related kinases CDK7, CDK8, and CDK19, the cell cycle kinase CDK1, and the cytoskeletal regulator MAK, all of which showed conserved activation across all species. Nutrient/metabolic sensing kinases showed a mixed response overall rather than a consistent trend, with mTOR in particular showing divergent regulation across the three coral species.

Given the strong and conserved response of canonical MAP kinases, we further interrogated the time course data to visualize ERK, JNK and p38 dynamics (Fig. 3C). This showed a steady increase in activity for all three coral species, starting as early as one hour post heat stress, indicating a clear role for MAP kinases in the early response to thermal stress. Orthogonal validation was carried out for phosphorylated-ERK in both *P. acuta* and *P. compressa*, showing a significant increase (*p* = 0.045 in *P. acuta*, *p* = 0.009 in *P. compressa*) in the ratio of phosphorylated-ERK to total ERK at the 4-hour time point, confirming the results observed in our phosphoproteomics dataset (Fig. 3D).

### Phosphorylation-driven remodeling reveals conserved biological responses to thermal stress

We next sought to determine whether thermal stress elicited conserved pathway-level responses across coral species. To do this, we performed rank-based gene set enrichment analysis (GSEA), using the log2 fold change of signal intensity between ambient and heat in experiment 2 (4 hours after initial heat treatment) of individual phosphorylation sites as the ranking metric and restricting the analysis to phosphoproteins detected across all three coral species with identifiable human orthologs. We then distinguished enriched Gene Ontology (GO) terms as abundance-driven, defined as significant in the global proteome either alone or together with the phosphoproteome, or phosphorylation-driven, defined as significant exclusively at the phosphoproteome level. This analysis revealed a striking difference between the two regulatory layers: abundance driven responses showed poor pathway enrichment, whereas most of the enriched pathways were observed within the phosphorylation dataset, suggesting substantial regulation of pathways by phosphorylation during this acute heat stress (Fig. 4A, S. table 5 and 6).

**Figure 4:**
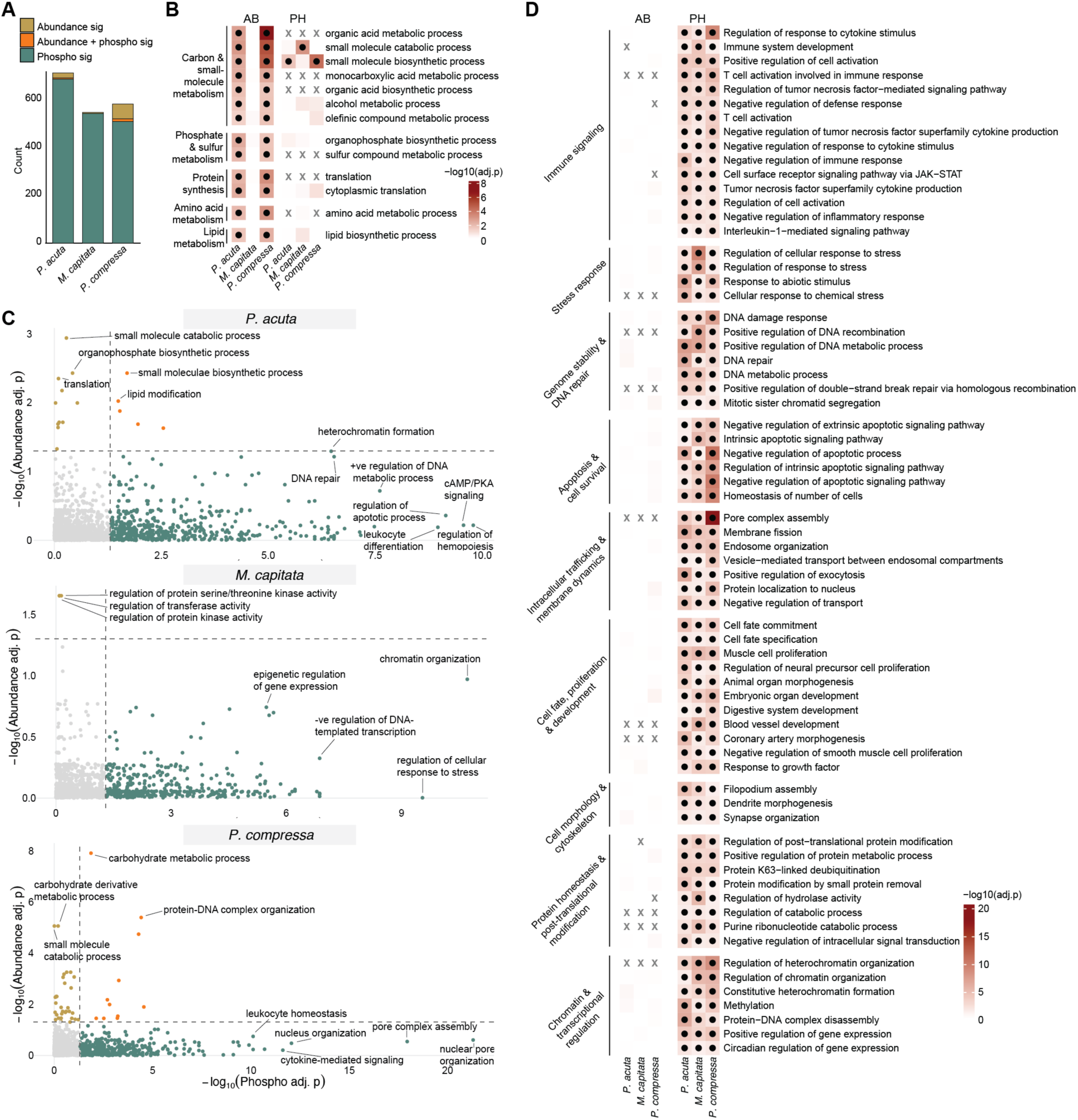
Phosphorylation-driven remodeling reveals conserved biological responses to thermal stress. **(A)** Proportion of abundance only-; abundance- and phosphorylation-; and phosphorylation only- driven changes in GO enriched pathways from the validation experiment (Exp. 2) across *Pocillopora acuta*, *Montipora capitata* and *Porites compressa*; **(B)** GO terms from enrichment of changes in protein levels that are significant in at least two species (i.e., either abundance only-driven or abundance- and phosphorylation-driven changes); **(C)** Scatter plots of the significance of phosphorylation-driven (X axis) and abundance-driven (Y axis) GO terms. **(D)** GO terms that are significant in all three species that are attributed to phosphorylation only-driven changes.

Among abundance-driven terms significant in at least two species, enriched processes grouped primarily into five broad functional categories: carbon and small-molecule metabolism, phosphate and sulfur metabolism, protein synthesis, amino acid metabolism, and lipid metabolism (Fig. 4B). Despite restricting the analysis to proteins conserved across all three species, the magnitude and significance of these abundance-level responses varied considerably among species (Fig. 4C), indicating divergence in proteome remodeling during thermal stress. In contrast, phosphorylation-driven responses shared across all three species revealed a highly conserved regulatory signature (Fig. 4D). These responses organized into nine functional programs encompassing immune signaling; stress responses; genome stability and DNA repair; apoptosis and cell survival; intracellular trafficking and membrane dynamics; cell fate, proliferation, and development; cell morphology and cytoskeletal regulation; protein homeostasis and post-translational regulation; and chromatin and transcriptional regulation. Taken together, our analysis provides a set of pathways that depict a conserved response to thermal stress, revealing common regulatory mechanisms underlying this response.

### Phosphorylation-driven immune signaling reveals extensive cellular remodeling during thermal stress

Our pathway enrichment analysis (Fig. 4D) revealed a strong enrichment of immune-related processes, including cytokine signaling, T cell activation, and regulation of inflammatory responses, with these changes occurring primarily at the phosphorylation rather than protein abundance level (Fig. 5A, fig. S5). To better understand the signaling architecture within the immune-related GO terms, proteins within enriched immune-related GO terms from Figure 4D were visualized as a network, with STRING-derived MCL-clustered functional associations represented as edges (Fig. 5A). Surrounding phosphosite halos depict heat-responsive phosphorylation events across the three coral species, with individual rings representing phosphosites and color indicating the direction and magnitude of phosphorylation changes relative to ambient conditions. The larger clusters were centered on intracellular signaling, such as cytokine- and stress-responsive MAPK-signaling, TNF/NF-kB signaling, and immune receptor signaling, consistent with broad regulation of inflammatory pathways during thermal stress. Overall, heat stress perturbed immune receptor signaling networks associated with non-self recognition, suggesting that thermal stress shifts the coral symbiosis from a tolerant, immunologically quiescent state toward one of heightened immune surveillance. This shift may reduce host tolerance of algal symbionts, offering a potential molecular explanation for symbiont expulsion during bleaching and revealing putative targets for enhancing coral resilience.

**Figure 5:**
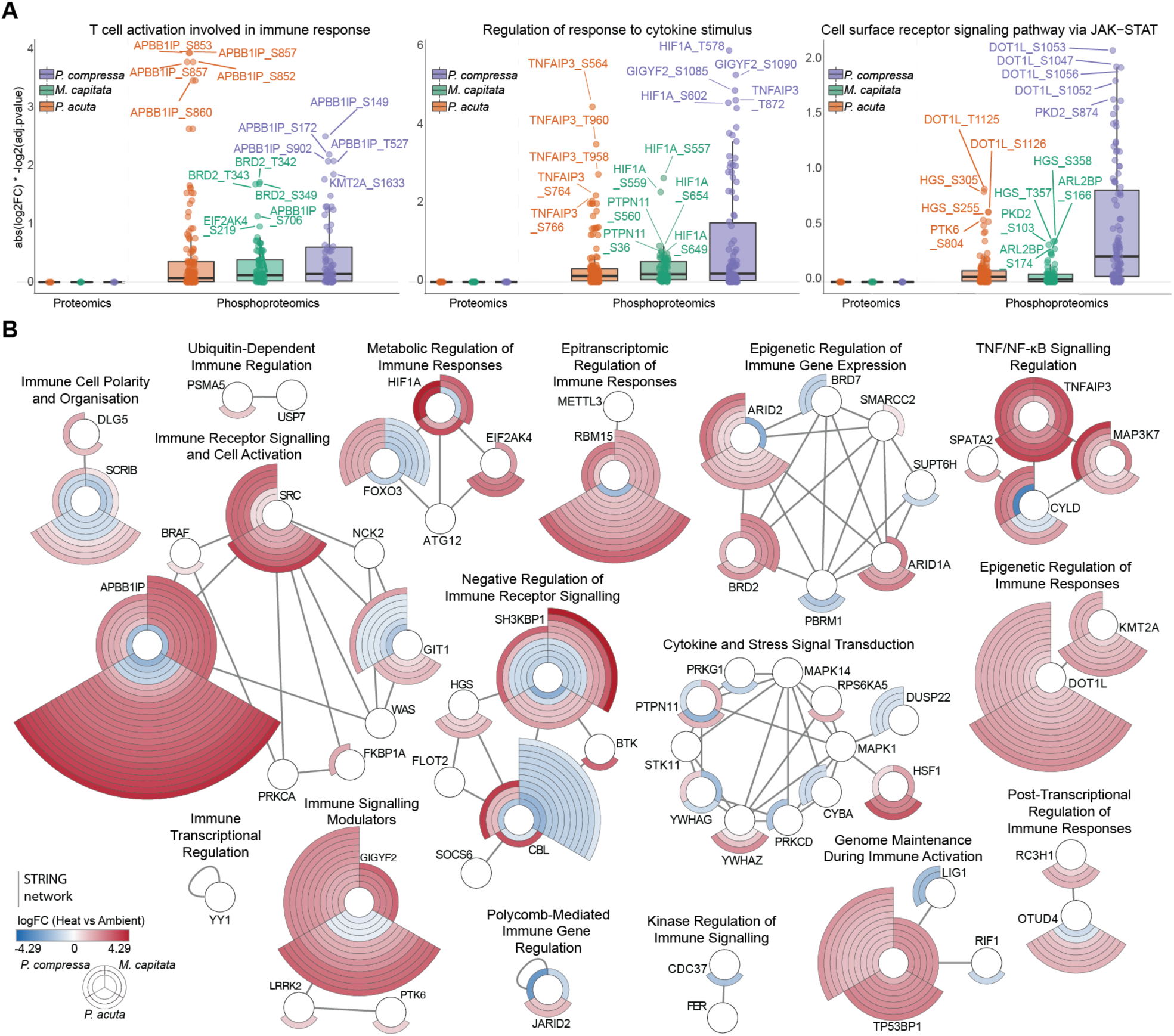
Phosphorylation-driven immune signaling reveals extensive cellular remodeling during thermal stress. **(A)** Relative abundances of proteins (left) and phosphorylation sites (right) from target GO terms (GO:0002286 - T cell activation in immune response; GO:0060759 - Regulation of response to cytokine stimulus; GO:0007259 - Cell surface receptor signaling pathway via JAK-STAT); **(B)** Induced subnetwork from STRING of proteins in enriched immune-related GO terms from cluster in Fig. 4D. Proteins are clustered using MCL clustering, only significant phosphorylation sites are displayed. Color depicts log2 fold change in heat versus ambient and halos, divided by species, depict individual phosphorylation sites detected along each protein, arranged from negative to positive.

Other clusters displayed epigenetic and transcriptional regulation, such as chromatin remodeling, Polycomb-mediated regulation, and histone methyltransferases, suggesting that heat stress may also modulate immune gene expression. There were clusters involved in regulation of post- transcriptional regulation, including RNA methylation and RNA stability, as well as clusters involved in autophagy and metabolic adaptations, DNA repair, cell polarity, and ubiquitin- dependent protein turnover, highlighting multiple layers through which thermal stress may remodel coral immune signaling. Interestingly, nearly all these individual phosphorylation sites remain functionally uncharacterized, yet many undergo substantial changes during heat stress, revealing a broad set of previously unexplored regulatory events and highlighting numerous opportunities for future investigation.

## DISCUSSION

This work aimed to characterize the phosphoproteome of three Hawaiian reef-building corals under acute thermal stress and apply bioinformatics techniques from model organisms to unravel signaling dynamics. Using novel approaches developed here to translate model systems knowledge networks to corals, we resolved structural conservation of kinases across kingdoms, a conserved MAPK stress response across species initiating a comprehensive signaling response that is difficult to detect in protein abundances. These data also confirm the primacy of the immune response during heat stress but suggest an earlier host-driven role in regulating this response before detectable strong oxidative stress responses.

Proteomics analysis resulted in the identification of between approximately 8000 and 11000 unique proteins in each species. This displays a marked increase in the current identifiable proteome from the literature and significantly increases the discoverable proteome. Additionally, we were able to optimize the phosphoproteomic pipeline for reef-building corals by flash-freezing, lyophilizing, and grinding up coral tissue, successfully navigating complexities such as the calcium carbonate skeleton, excessive salt and lipid content, and the pigments from host and symbiont. Our improved approach eliminated the need to separate host tissue from symbionts, allowing for less loss. After optimization, we were able to identify between approximately 19000 and 35000 unique phosphopeptides across the three species within 4 hours of heat exposure, numbers that are state-of-the-art even in the field of biomedicine (*39*). Moreover, this work demonstrates that phosphoproteomics captured signaling level regulation during acute heat stress that was largely invisible to abundance-based proteomics alone, underscoring the importance of phosphoproteomics for resolving the signaling pathways underlying the coral heat stress response. Additionally, we found over 70% of human kinases had a matching ortholog in corals, permitting the application of prior knowledge from human studies to characterize dysregulated kinases and pathways in corals.

Despite representing distinct scleractinian families, *P. acuta*, *M. capitata*, and *P. compressa* exhibited a conserved signaling response to heat stress, suggesting this response extends beyond a single lineage. A prominent aspect of this shared signaling was the activation of canonical MAP kinases in the ERK, JNK, and p38 pathways. MAP kinase signaling is known to play a vital role in early cellular responses to stress, relaying information from sensors to enact cellular responses, a process conserved in eukaryotes(*40*). Even during osmotic stress in yeast, MAP kinases coordinate stress responses temporally, spatially, and mechanistically, enabling adaptation to harsh environments (*41*). Within the coral literature, evidence has shown increased activity of MAPKs p38 and MAPK3K1 under thermal stress (*42*). Additionally, corals have been shown to activate JNK to repress stress-induced reactive oxygen species (ROS) accumulation (*43*). The accumulation of ROS has been clearly demonstrated to play a central role in temperature- and light-mediated bleaching (*44*); however it remains unclear whether ROS precedes or follows MAPK activation. In our data, the enrichment of ROS related terms centers on MAPK enrichment (Fig. 3), and although these processes are interrelated, our data cannot distinguish the order of activation.

In addition to MAPK activation, one of the most prominent conserved features of the coral response to thermal stress was the phosphorylation-driven remodeling of immune-related pathways. Thermal stress is being increasingly recognized to induce both rapid and dynamic remodeling of immune function in cnidarians, with evidence for modulation of immune gene expression and immune cell activity (*45–50*). Phagocytosis and pinocytosis assays can be used as a proxy for immune cell activity, and both *Nematostella vectensis* and Aiptasia display an increased phagocytic activity under heat stress (*47*). A recent study built further on this using transcriptomics to demonstrate stress-induced activation of conserved innate immune genes in *N. vectensis*, Aiptasia, and the stony coral *Stylophora pistillata,* during heat stress (*45*).

Transcriptomics responses represent one level of immune regulation and may happen downstream of the initial phosphorylation-driven signaling stimulus that has initiated the response.

Our data revealed phosphoregulation of an immune signaling network that merges components classically associated with both innate and adaptive immunity in vertebrates. Several of the identified proteins act as proximal signal transducers immediately downstream of pattern recognition receptors (i.e., innate immunity) and antigen receptors (i.e., adaptive immunity), including SRC, NCK2, WAS, BTK, CBL, SH3KBP1, GIT1, and PTPN11 (Fig. 5). Although corals lack an adaptive immune system, and none of the canonical T or B cell receptor machinery is present in cnidarian genomes, we interpret these proteins through the lens of deep homology, in which ancient receptor signaling and cytoskeletal remodeling components are independently reused across evolutionary contexts (*51*). In mammals, SRC family kinases initiate signaling downstream of the T cell receptor, B cell receptor, and Fc receptors by phosphorylating receptor-associated immunoreceptor tyrosine activation motifs (ITAMs), while NCK2 couples to CD3ε to regulate T cell receptor internalization (*52*). WAS drives the actin remodeling required for immunological synapse assembly and sustained T cell receptor signaling (*53*). Bruton’s tyrosine kinase (BTK), best known for its role in B cell receptor signaling and B cell development, also acts in the innate immune system, downstream of Toll-like and Fc receptors to promote cytokine production in a variety of cell types (*54*), while CBL functions to negatively regulate receptor kinases for proteasomal degradation (*55–57*). Transforming growth factor beta (TGFβ) binding to TGFβ receptors activates PRKCA, which promotes SMAD activation (*58*) and the production of pro-inflammatory cytokines (*59*). SMAD proteins aid in the cellular response to oxidative stress (*60*), and there is an enrichment of SMAD motifs under heat stress in Aiptaisa (*61*). This suggests that coral immune receptor signaling possibly relies on an ancient, conserved toolkit that predates the evolutionary split between innate and adaptive immunity in vertebrates.

Downstream of pathogen recognition (i.e., Toll-like) and antigen receptors lies NF-κB (Fig. 5). BTK and PTPN11 both promote NF-κB activation and cytokine or interferon production downstream of Toll-like receptors (*62*, *63*), while CBL, OTUD4, and USP7 act as regulators, with OTUD4 suppressing and USP7 enhancing NF-κB activity through deubiquitination of MyD88 (*64*, *65*). Consistent with this, NF-κB is reported to be upregulated during heat stress across coral species (*66–68*), and Toll-like receptor signaling to NF-κB has been described in the reef-building coral *Orbicella faveolata* (*69*). This suggests a coordinated activation of this proximal signaling layer alongside regulation of its downstream convergence point, NF-κB, presenting a unified picture in which signaling components classically divided into innate and adaptive immunity in vertebrates jointly regulate the coral response to heat stress.

Taken together, our findings are consistent with a model in which the earliest determinants of coral bleaching are set by host signaling decisions within hours of acute thermal stress, reframing bleaching as an active, host-initiated response rather than purely stress-driven physiological collapse. Across three evolutionarily distinct reef-building corals, acute heat stress induced a conserved phosphosignaling program characterized by MAPK activation and the regulation of immune receptor-proximal signaling machinery associated with non-self recognition, suggesting that thermal stress may shift the coral host away from symbiont tolerance and toward conditions permissive for symbiosis rejection. Because only a minority of coral cells are thought to harbor algal symbionts, both within the gastrodermis specifically (*70*) and across the whole polyp (*71*), a substantial component of this signal plausibly originates from aposymbiotic host cells (*72*). Although this does not exclude the possibility of a disproportionate contribution from symbiont-containing cells themselves, our data point to a rapid, tissue-wide host signaling program that may help determine whether symbiosis is maintained or dismantled. By revealing this otherwise invisible regulatory layer, our study establishes phosphoproteomics as a powerful approach for resolving the earliest molecular events of the coral thermal stress response and identifies conserved host signaling pathways as candidate targets for understanding, and potentially enhancing, coral resilience to a warming ocean.

## MATERIALS AND METHODS

### Permits for coral collection

All corals were collected under SAP 2024-45 and SAP 2025-26 from the Hawaiʻi Department of Land and Natural Resources - Division of Aquatic Resources to HIMB.

### Coral collection and experimental set-up

We conducted two separate acute heat stress time course experiments: a main experiment and a subsequent validation. Fragments (∼10cm total branch linear extension) from each of 3 colonies of *Montipora capitata*, *Porites compressa* and *Pocillopora acuta* were collected from the reef flat at the Hawaiʻi Institute of Marine Biology (HIMB; 21.4309, -157.7873). For the first experiment, we collected corals on the afternoon of 2 January, 2024. Corals were split into replicate ∼5cm branches, immediately mounted to aragonite plugs and haphazardly assigned to two tanks (ambient control and a heat ramp at HIMB). In total, there were single fragments of each colony (n=3) of each species (n=3) in each treatment (n=2). Tanks under ambient light were maintained at ambient temperature (24°C) until 1000h the next morning, at which point the heated treatment was ramped to 32°C over one hour (Fig. 1A; peaking at 1100h) and maintained for the duration of the experiment (1400h). The control tank was maintained at 24°C over the same time course.

The second experiment was conducted in the same framework, with new collections of 3 novel colonies from each species from the same location on 4 April, 2025. The timeline used to validate data was identical to the initial trial.

### Coral sample harvest

In experiment 1, corals in both treatments were sampled before the initiation of heat stress (“Pre-Heat”; 1000h; Fig. 1A) and after 4h (1400h). Heated corals were also sampled at 1h (1100h), coinciding with treatments reaching peak temperatures, and at 2h (1200h). At each time point, every fragment was sampled in triplicate using a 5mm dermal curette, which was immediately flash frozen in liquid nitrogen.

In the second experiment, novel corals were used with the same harvest design, mounting, tank acclimation and experimental heat ramp, but were sampled only at 4h (1400h) by flash freezing entire fragments immediately in liquid nitrogen.

### Protein extraction

For experiment 1, samples were resuspended in 500 µL ice-cold 6M GuHCl lysis buffer and homogenized with 0.3 g glass beads and 20 µL protease inhibitor (PI; Sigma-Aldrich) by bead beating at maximum speed for 3 min. The homogenate was transferred to a fresh tube and centrifuged at 1,400 × g for 5 min to separate host material (supernatant) from symbiont cells (pellet). The host supernatant was transferred to a low-binding tube and kept on ice. β-mercaptoethanol (5 µL) was added, followed by incubation at 85 °C for 10 min. Samples were extracted with 1 mL ethyl acetate, vortexed for 1 min, and centrifuged at 10,000 × g for 1 min. The upper organic phase was removed, and the ethyl acetate extraction was repeated twice. The resulting lower aqueous phase was retained for subsequent analysis. This aqueous phase was centrifuged at 20000 x g for 10 mins and 200 µL of the supernatant (excluding any precipitated skeleton) was transferred to a sterile Protein LoBind tube. A chloroform-methanol precipitation was then performed with the step-wise addition of ice-cold methanol (4 x volume -800 µL), chloroform (1 x volume - 200 µL), and water (3 x volume - 600 µL) with thorough vortexing at each step. Samples were centrifuged at 14000 x g for 10 mins, and the upper and lower aqueous phases were poured away together leaving the protein pellet behind. The protein pellet was then washed twice with ice cold methanol, centrifuging at 20000 x g each time. The pellets were air dried before being resuspended in an 8M urea lysis buffer.

For experiment 2, a more efficient pipeline was implemented. Flash frozen coral fragments were crushed by pestle and mortar under liquid nitrogen before being lyophilized until dryness. This rubble-like material was crushed to a fine powder using a pestle and mortar once more. 400 µL of 8M Urea lysis buffer was added and samples were vortexed to combine, before being centrifuged at 20000 x g for 10 mins at 4 and 200 µL of supernatant transferred to a sterile 2 mL Protein LoBind tube. A chloroform-methanol precipitation was then performed as mentioned in the previous paragraph.

### Sample processing for proteomics and phosphoproteomics

Lysed samples were sonicated using a probe sonicator 5x for 10 seconds On and 10 seconds Off at 10% amplitude, and protein was quantified using a bicinchoninic acid assay (BCA). Approximately 200 µg of protein for each sample was used for further processing, starting with reduction and alkylation using tris-(2-carboxyethyl) (TCEP) (10 mM final) and 2-chloroacetamide (40 mM final) for 30 minutes each at 24°C with shaking at 1200 rpm. Before protein digestion, the 8 M Urea was diluted 5-fold with 100 mM Tris-HCl (pH 8) to permit the activity of the proteolytic enzyme trypsin. Trypsin (Promega) and Lys-C (Wako) was added at a 1:100 (wt/wt) and 1:50 (wt/wt) enzyme-substrate ratio, respectively, and placed in a thermomixer at 24°C overnight (16 h) with shaking at 1200 rpm. Following digestion, 10% trifluoroacetic acid (TFA) was added to each sample to a final pH of approximately 2-3. Samples were then desalted using a vacuum manifold with Oro-Flex I Polypropylene filter plate w/10 m PE frit 96 well plate (Orochem) with 7 mg HLB Resin (Oasis) for 100 ug of peptide. Each well with resin was activated with 200 µL of 100% acetonitrile (ACN), then equilibrated with 3 x 200 µL of 0.1% TFA. After sample loading, the wells were washed with 3 x 200 µL of 0.1% TFA, and samples were eluted with 2 x 80 µL 50%ACN/0.25% formic acid (FA). Approximately 10% of the desalted peptides were separated for global abundance proteomics analysis and the remaining 90% were used for phosphopeptide enrichment. Both the fractions were then dried by vacuum centrifugation in a SpeedVac (Labconco).

For phosphopeptide enrichment, Ti-IMAC beads (Resyn Biosciences) were aliquoted in a 1:2 (w/w) peptide:beads ratio and equilibrated three times with binding buffer (0.1 M glycolic acid in 80% acetonitrile (ACN), 5% TFA). Dried peptide samples were resuspended in 200 µL of binding buffer, added to the equilibrated beads, and incubated for 30 min at 24 °C, 1200 rpm. The unbound fraction was discarded, and beads were washed sequentially once each with 100 µL of the binding buffer, wash buffer 1 (60% ACN, 1% TFA, 200 mM NaCl), wash buffer 2 (60% ACN, 1% TFA), and finally with LC-MS grade water. Enriched phosphopeptides were eluted twice by incubating the beads with 150 µL of 1% (v/v) ammonium hydroxide (Sigma) in LC-MS grade water (Fisher Scientific) for 10 min at 24 °C, 1200 rpm. The eluted peptides were each transferred to a new protein LoBind tube containing 50 µL of 10% (v/v) formic acid in LC-MS grade water. Both eluates were pooled and dried followed by plate-based desalting as explained previously and were further resuspended in 0.1% formic acid. A volume corresponding to approximately 500 ng of peptides was injected into the timsTOF HT mass spectrometer (Bruker Daltonics).

### Mass spectrometry proteomics acquisition

Dried peptides were resuspended in 0.1% (v/v) FA in MS grade water (Fisher) and analyzed on a Bruker timsTOF HT mass spectrometer, paired with a ThermoFisher Vanquish Neo UHPLC system. Mobile phase A consisted of 0.1% (v/v) FA in MS grade water (Fisher), and mobile phase B consisted of 0.1% (v/v) FA in 100% MS grade Acetonitrile (Fisher). The LC was operated in trap-and-elute mode, where the peptides were first trapped onto a PepMap Neo Trap column (5 mm, 100 Å pore size, 5 µm particle size) and then reversed-phase separated using gradients mentioned below on an Aurora Elite C18 reverse phase column (15 cm, 100 Å pore size, 1.5 µm particle size for captive spray, IonOptiks), kept at 50°C using a column oven for Bruker Captive Spray source (Sonation Lab Solutions), and ionized in a CaptiveSpray source (Bruker Daltonics) at 1700 V. For global abundance proteome analysis, the %B gradient used was: 5% to 35% over 37 min at 0.3 µL/min, then to 45% in the next 4 mins, and 60% in the next 1 min, followed by an increase to 95% B over 3 min. For phosphoproteome analysis, the %B gradient used was 3% to 16% over 26 min, 30% in the next 11.5 min, 45% in the next 4 min, and 60% B for 1 followed by an increase to 95% B over 3 min at 0.3 µL/min flow rate.

For the global abundance data analysis, equal-size windows of 25 Da were designed with an overlap of 1 Da to maximize the precursors ion coverage for further MS/MS. Ions were accumulated and separated in the TIMS analyzer over a mobility range of 0.60–1.60 V·s/cm² using 100 ms accumulation and ramp times. DIA acquisition comprised one MS1 scan followed by eleven dia-PASEF scans covering a mass range of 384.4–1305.4 m/z and an ion mobility range of 0.77–1.41 1/K0. Collision energies were linearly ramped according to ion mobility from 20 eV at 1/K0 = 0.71 to 59 eV at 1/K0 = 1.41. The resulting cycle time was approximately 1.27 s. For phosphoproteome analysis, the raw data was acquired in dia-PASEF mode with variable isolation window widths in the m/z vs ion mobility plane. For dia-PASEF acquisition, ions were accumulated and separated in the TIMS device over a mobility range of 0.60–1.60 Vs/cm². MS/MS spectra were acquired across a mobility range of 0.67–1.33 1/K0 and a mass range of 307.2–1292.2 m/z. Collision energies were linearly ramped as a function of ion mobility from 20 eV at 1/K0 = 0.71 to 59 eV at 1/K0 = 1.41. The estimated mean cycle time was 1.27 s. For the global abundance data analysis, equal-size windows of 25 Da were designed with an overlap of 1 Da to maximize the precursors ion coverage for further MS/MS.

### Mass spectrometry proteomics data search and quantitative analysis

The raw files were processed with Spectronaut (Biognosys, [Update 20.2.250922.92449]) with the directDIA+ (Deep) search algorithm, its *in silico* derived DIA analysis. Carbamidomethylation (cysteine) was set as a fixed modification for database search. Acetylation (protein N-term), oxidation (methionine), and phosphorylation (serine, threonine, tyrosine; only for phosphoproteomics dataset) were set as variable modifications. Reviewed human protein sequences (downloaded from UniProt, October 6, 2023) were used for spectral matching. The false discovery rates for the PSM, peptide, and protein groups were set to 0.01, and the minimum localization threshold for PTM was set to zero. For MS2-level area-based quantification, the cross-run normalization option was unchecked (normalization was performed later using MSstats), and the probability cutoff was set to zero for the PTM localization. Quantitative analysis was performed in the R statistical programming language (v.4.4.1).

Initial quality control analyses, including inter-run clustering, correlations, principal component analysis (PCA), peptide and protein counts, and intensities were completed in R. Statistical analysis of phosphorylation and protein abundance changes between exposed and control samples were computed using the R package MSstats (4.12.1). For protein abundance, all peptides mapping to the same proteins were summarized together using the Tukey’s Median Polish approach. For phosphoproteomics, all peptides containing the same set of phosphorylated sites were summarized together into phosphorylation site groups using Tukey’s Median Polish. For both phosphopeptide and protein abundance MSstats pipelines, MSstats performs normalization by median equalization, imputation was turned Off (set to FALSE), and statistical tests of differences in intensity between conditions were calculated using default settings in MSstats. Specifically, MSstats calculates log2 fold changes as the ratio of averaged (across replicates) protein or phosphopeptide intensities between conditions, uses a Student’s t-test for p-value calculation and the Benjamini-Hochberg method of FDR estimation to adjust p-values.

### Kinase homology and conservation analysis

To better understand the functional implications of heat exposure on reef-building corals, we used OrthoFinder to match coral proteins to their human orthologs (Supplementary Table 1) (*34*). When multiple coral proteins matched to a single human gene, we used sequence similarity to select the closest match within orthogroups. For human kinases with a corresponding coral protein, we performed Needleman-Wunsch global sequence alignments to quantify sequence similarity. In parallel, we used AlphaFold2 to predict structures for all matched human and putative coral kinases and calculated TM-scores to quantify structural similarity based on the length-normalized spatial distances between aligned residues, with higher scores indicating greater overall structural conservation.

### Kinase activity inference

The Kinase Library was used to infer kinase activity (*37*). Phosphoacceptor sites (S, T, Y) that were experimentally detected were extracted from the representative coral genome along with the log2FC value, with the surrounding 6 amino acids (3 either side). The Kinase Library is species agnostic, meaning no data had to be filtered for matches to human orthologs. The differential enrichment tables were downloaded for each species, and a heatmap was created using the ComplexHeatmap package (v2.26.1). The resulting kinases were manually grouped into broad functional groups using literature review, and clustered within groups using the ward.D2 metric. The broader functional groups were then subject to a pathway enrichment analysis using the clusterProfiler package (v4.18.4), using the C5 Gene Ontology collection of the Molecular Signatures Database (MSigDB), accessed through the msigdbr R package (v26.1.0). The top 5 terms were displayed per functional grouping.

### Gene Enrichment analysis

Gene Ontology (GO) enrichment analysis was performed using the rank-based method, GO_MWU (*73*), which uses the Mann-Whitney U test to determine if genes annotated to a given GO category are non-randomly distributed within a genome-wide ranked list. Only coral genes with a matching human ortholog and were experimentally detected in all three species were considered. GO annotations were obtained from the C5 (Gene Ontology) collection of the Molecular Signatures Database (MSigDB), accessed through the msigdbr R package (v26.1.0), restricted to human gene symbols and excluding the Human Phenotype Ontology (HPO) sub-collection. The GO hierarchy was obtained from go.obo (Gene Ontology Consortium, release date 2026-06-28 (*74*, *75*)). Analyses were run separately for each species on Biological Processes domain separately for the proteomics and phosphoproteomics data, using these parameters: GO categories comprising >10% of all annotated genes or <5 genes were excluded; hierarchical clustering of gene-sharing categories used a cut height of 0.25. For abundance driven changes all terms significant in the proteomics data were considered, regardless of their status in the phosphoproteomics data. For visualization purposes, all terms significant in 2 or more species in the proteomics enrichment were considered, matching in the data from the phosphoproteomics enrichment. A heatmap was made in R using the ComplexHeatmap package (v2.26.1). For the phosphorylation driven changes, only terms that were significantly enriched in the phosphoproteomics data were considered. Due to the high number of significantly enriched terms, only those significant in all three species were visualized, matching in the data from the proteomics enrichment. Both heatmaps were subsequently clustered into broader functional groups manually in Adobe Illustrator based on prior knowledge and literature searches.

### Network analysis and visualization

All proteins that were assigned to the GO terms classified as immune-related were extracted and clustered using the STRING (v12.0) open-source database using the web interface. MCL-clustering was used with the default parameters and the individual nodes were exported. Cytoscape (v3.10.4) was used to visualize the clusters and the phosphorylation data for each species was overlaid. The phosphorylation sites were ordered from lowest to highest logFC and only significant sites were used for this visualization.

## SUPPLEMENTARY FIGURES

**Figure S1:**
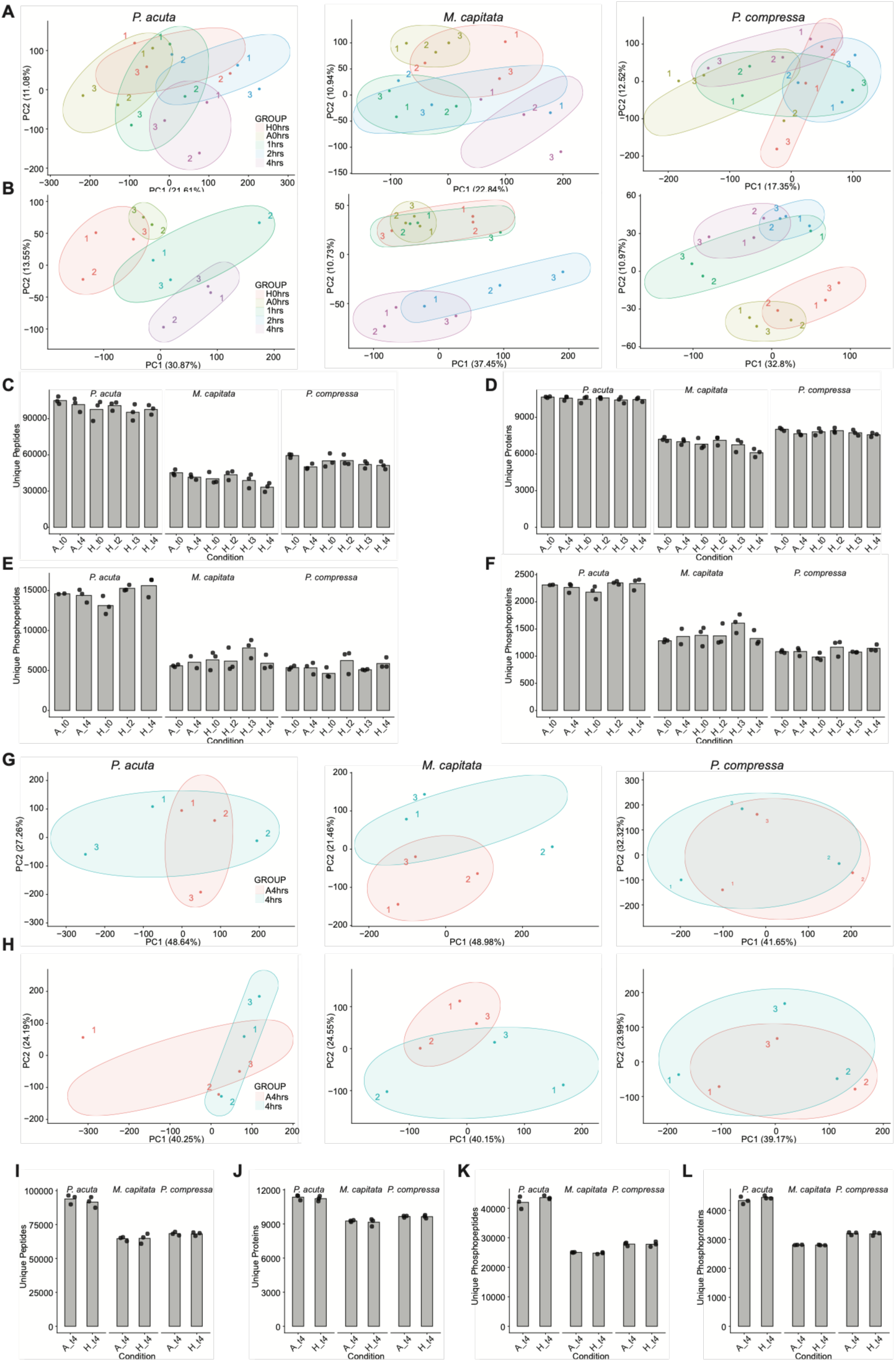
**(A)** Principal component analysis of the proteomic profiles of *Pocillopora acuta*, *Montipora capitata*, and *Porites compressa* in experiment 1; **(B)** Principal component analysis of the phosphoproteomic profiles of *Pocillopora acuta*, *Montipora capitata*, and *Porites compressa* in experiment 1; **(C)** Unique peptides detected in experiment 1; **(D)** Unique proteins detected in experiment 1; **E**. Unique phosphopeptides detected in experiment 1; **(F)** Unique phosphoproteins detected in experiment 1; **(G)** Principal component analysis of the proteomic profiles of *Pocillopora acuta*, *Montipora capitata*, and *Porites compressa* in experiment 2; **(H)** Principal component analysis of the phosphoproteomic profiles of *Pocillopora acuta*, *Montipora capitata*, and *Porites compressa* in experiment 2; **(I)** Unique peptides detected in experiment 2; **(J)** Unique proteins detected in experiment 2; **(K)** Unique phosphopeptides detected in experiment 2; **(L)** Unique phosphoproteins detected in experiment 2.

**Figure S2:**
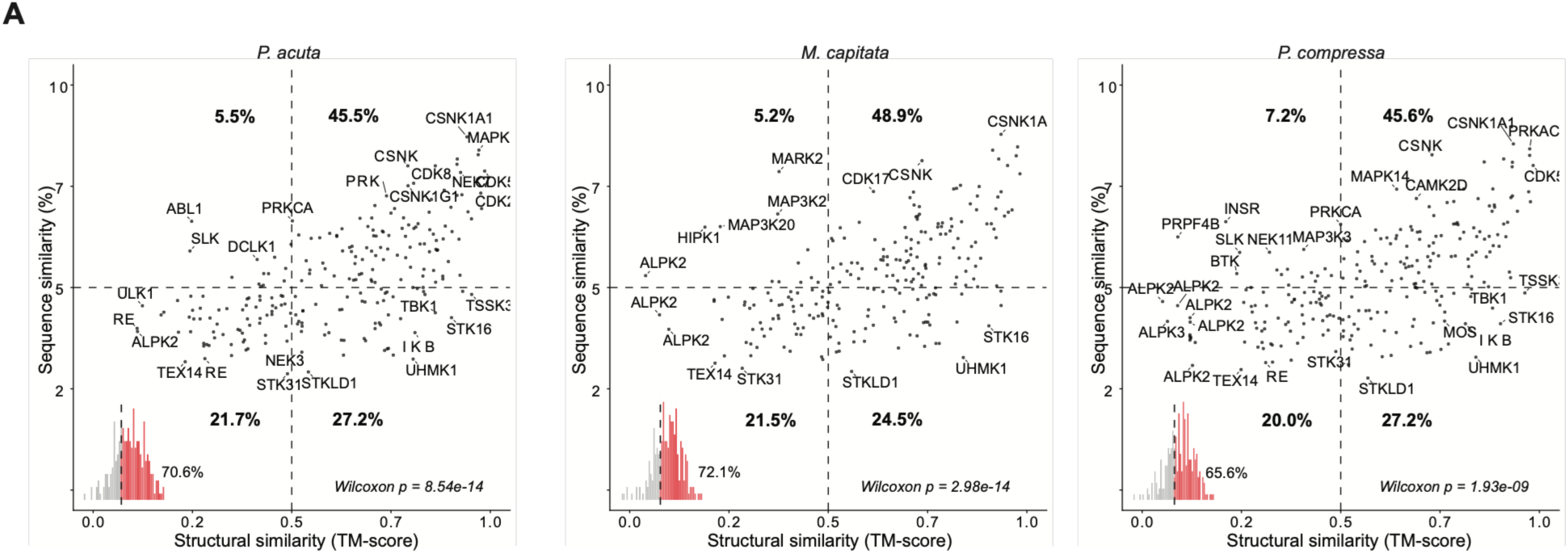
**(A)** The correlation of structural and sequence similarity of identified putative coral kinases, with both a match to a human kinase, and an active kinase domain, split by species.

**Figure S3:**
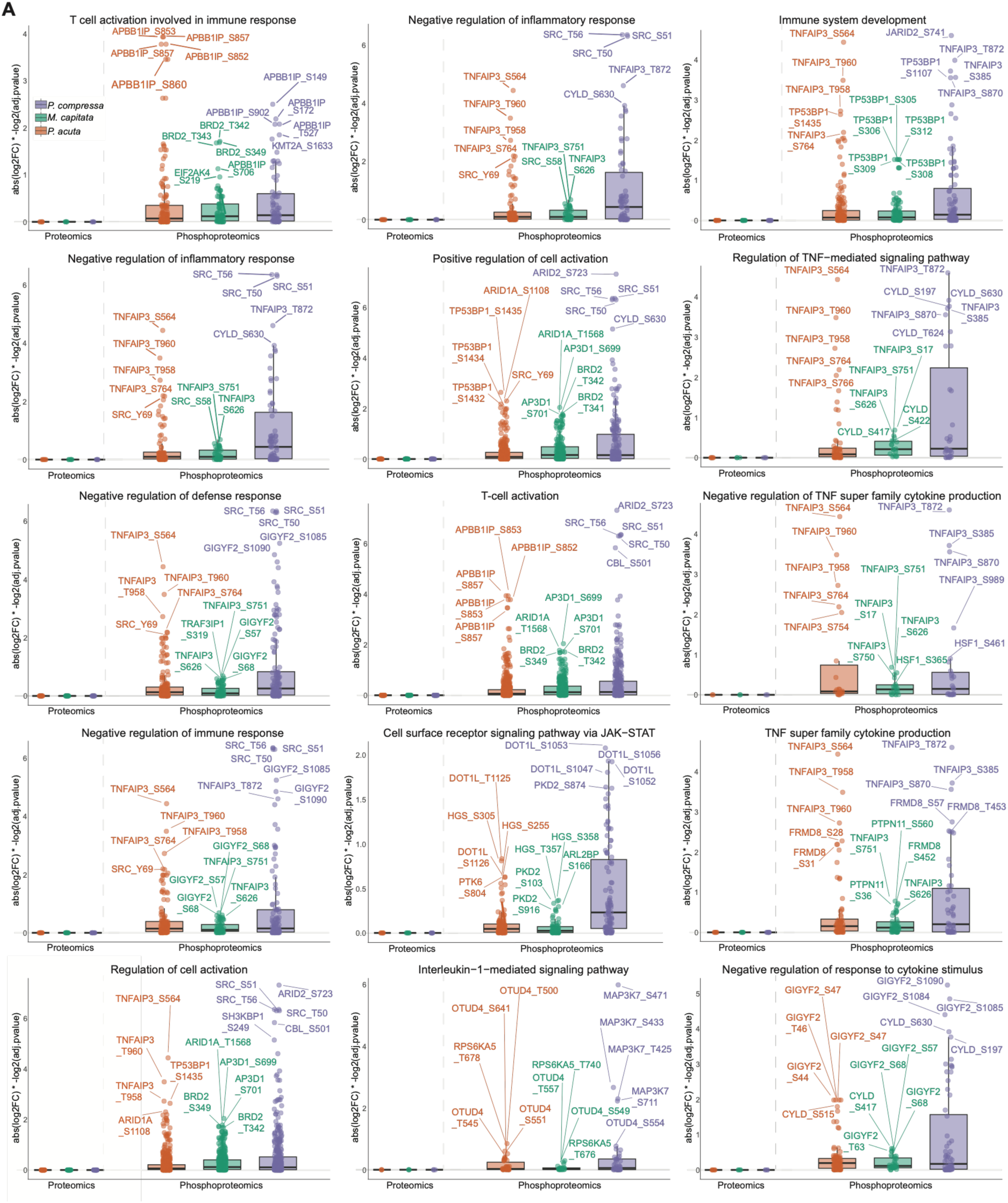
**(A)** Relative abundances of protein abundances and phosphorylation sites from target GO terms related to immune response detected in the GO enrichment analysis.

## SUPPLEMENTARY TABLES

**Table S1.** Orthofinder analysis of coral proteins matched to human proteins, with highest coral sequence similarity used to match each human protein.

**Table S2.** Expanded orthofinder analysis of all possible matches, including matches across the three coral species.

**Table S3.** The TM structural similarity and sequence similarity scores for kinase matches between human kinases and putative coral kinases.

**Table S4.** Differential enrichment scores for serine and threonine kinases for *Pocillopora acuta*, *Montipora capitata*, and *Porites compressa* at 4 hours of heat treatment.

**Table S5.** Biological Processes GO_MWU proteomics and phosphoproteomics enrichment data for *Pocillopora acuta*, *Montipora capitata*, and *Porites compressa* at 4 hours post heat treatment.

**Table S6. (On Mendeley Data)** Biological processes heats file for proteomics and phosphoproteomics enrichment data for *Pocillopora acuta*, *Montipora capitata*, and *Porites compressa* at 4 hours post heat treatment. Due to size, it will be made available on Mendeley Data. (https://data.mendeley.com/preview/3rgj27z7ms?a=81023b11-158e-4b88-8e3d-44790a2d337c)

## SUPPLEMENTARY DATA

**Data S1. (On Mendeley Data)** All Alphfold2 structures for human kinases and putative coral kinases. Due to size, it will be made available on Mendeley Data. (https://data.mendeley.com/preview/3rgj27z7ms?a=81023b11-158e-4b88-8e3d-44790a2d337c)

## ACKNOWLEDGEMENTS

We would like to acknowledge the lab of Heather D. Maynard at UCLA for allowing us to use their lyophilizer.

## FUNDING

Walder Family Foundation (CD and GT)

UCLA AIDS Institute (MB)

James B. Pendleton Charitable Trust (MB)

McCarthy Family Foundation (MB)

National Institutes of Health (NIH-NIGMS R35GM160071 to MB)

Cellular and Molecular Biology T32 at UCLA (NIH-NIGMS T32GM145388 to DMW)

Microbial Pathogenesis Training Grant at UCLA (MPTG NIH-NIAID T32AI007323 to DMW)

Undergraduate Research Scholars Program Grant funded by the Van Trees at UCLA (USRP, 2026-2027 to SAC).

## AUTHOR CONTRIBUTIONS

Conceptualization: IAA, CD, MB

Experimental performance: CD, GT, IAA, EM, SR, TP

Sample processing: IAA, JN, DMW, SAC

Data analysis: IAA, MB, CD, KA

Manuscript writing original draft: IAA, MB, CD, GT, EM

Manuscript writing review: IAA, MB, CD, GT, EM, SR, DMW, SAC, JN, KA, TP

## DATA, CODE, AND MATERIALS AVAILABILITY

Data are available via ProteomeXchange PRIDE with identifier PXD084041, token cRQOT6k4spRc. Supplementary table 6 and Supplementary Data 1 are available on Mendeley Data (https://data.mendeley.com/preview/3rgj27z7ms?a=81023b11-158e-4b88-8e3d-44790a2d337c).

## SUPPLEMENTARY MATERIALS

Materials and Methods

Figs. S1 to S3

Tables S1 to S6 Data S1

